# Molecular basis for Ras suppressor-1 recruitment to focal adhesions and stabilization of consensus adhesome complex

**DOI:** 10.1101/2021.02.19.432017

**Authors:** Koichi Fukuda, Fan Lu, Jun Qin

## Abstract

Ras suppressor-1 (Rsu-1) is a leucine-rich repeat (LRR)-containing protein that is crucial for regulating fundamental cell adhesion processes and tumor development. Rsu-1 interacts with a zinc-finger type multi LIM domain-containing adaptor protein PINCH-1 involved in the integrin-mediated consensus adhesome but not with highly homologous isoform PINCH-2. However, the structural basis for such specific interaction and regulatory mechanism remains unclear. Here, we determined the crystal structures of Rsu-1 and its complex with the PINCH-1 LIM4-5 domains. Rsu-1 displays an arc-shaped solenoid architecture with eight LRRs shielded by the N- and C-terminal capping modules. We show that a large conserved concave surface of the Rsu-1 LRR domain recognizes the PINCH-1 LIM5 domain, and that the C-terminal non-LIM region of PINCH-2 but not PINCH-1 sterically disfavors the Rsu-1 binding. We further show that Rsu-1 can be assembled, via PINCH-1-binding, into a tight hetero-pentamer complex comprising Rsu-1, PINCH-1, ILK, Parvin, and Kindlin-2 that constitute a major consensus integrin adhesome crucial for focal adhesion assembly. Consistently, our mutagenesis and cell biological data consolidate the significance of the Rsu-1/PINCH-1 interaction in focal adhesion assembly and cell spreading. Our results provide a crucial molecular insight into Rsu-1-mediated cell adhesion with implication on how it may regulate tumorigenic growth.

## Introduction

The adhesion of cell-extracellular matrix (ECM) is one of the essential mechano-chemical process for the life of multicellular organisms. The ability of cell-ECM adhesion is primarily mediated by cell surface receptors, integrins (1), and their adhesive properties significantly impact on a large number of fundamental physiological processes such as development, tissue organization, and proper functionality of cells (2). Integrins engage multi-layers of intracellular protein-protein interactions to connect ECM with the actin cytoskeleton, thereby transmitting their mechano-chemical signals into downstream effectors (3). The integrin-mediated adhesion complex (integrin adhesome) (4) was shown to comprise 232 components (2,4) within four distinct cluster modules in the core cell adhesion machinery (2,5). Dysregulated function of integrin adhesome components is critically linked to diseases such as cancer progression and metastasis that comprise a multi-step process (6). Deconvolution of the mechanism for the assembly and regulation of integrin adhesion complex could thus fill fundamental gaps in the unresolved interaction network to advance potential therapeutic intervention.

Rsu-1 (Ras suppressor-1) is one of the as-yet-understudied proteins that are enriched at the integrin adhesion complex (5). Rsu-1 is evolutionarily conserved throughout mammalian and invertebrate development (7-9). Previous genetic study demonstrated that a null mutation of Rsu-1 resulted in wing blistering in *Drosophila* that supports its crucial role in an integrin-dependent cell adhesion process (8). Rsu-1 does not have catalytic activity but its primary sequence analysis indicates a leucine-rich repeat (LRR)-containing protein (7) that implicates for protein-binding functions (10,11). At the molecular level, Rsu-1 interacts with a zinc-finger-type five LIM domains-containing adaptor protein PINCH-1 (8,12) that plays a crucial role in cell shape, migration, and survival (13). PINCH forms a complex with integrin linked kinase (ILK) pseudokinase, which in turn binds Parvin, constituting the heterotrimer IPP (ILK-PINCH-Parvin) complex (14). The IPP complex is a crucial central hub at the downstream of integrin network (5,15), and Rsu-1 cooperates with the IPP complex for cell adhesion and spreading (16). Notably, a previous study has identified that Rsu-1 has a role in impaired migration of MCF10A breast epithelial cells and has been implicated in upregulation of basal high-grade breast tumors (17). Indeed, those impaired migration phenotypes were shown to involve many focal adhesion proteins including PINCH-1 in the β-1 integrin network (17). However, the molecular and regulatory mechanisms by which the Rsu-1-PINCH axis governs cellular motility and signaling events remain unclear.

Rsu-1 was originally identified as a suppressor of Ras-dependent oncogenic transformation (7). Since Ras proteins regulate a variety of signal transduction responsible for cancer progression (18), several Ras-dependent pathways were proposed to link with Rsu-1 such as the Ras-MAPK pathway during the development of oncogenesis. Overexpression of Rsu-1 in epidermal growth factor-stimulated model cells with NIH3T3 and PC12 was shown to result in the inhibition of c-Jun N-terminal Kinase (JNK) and activation of ERK in relation to the Ras-MAPK pathway (19). Interestingly, the interaction between Rsu-1 and PINCH-1 was linked to the stabilization of the IPP complex (20), the regulation of the JNK signaling, development, and maintenance of organisms (8,9,12,21). A genome-wide gene expression study also revealed that Rsu-1 was one of the 93 LRR genes with elevated expression in the immune cluster from the tissue microarray (22). These studies imply potential unique scaffolding mechanisms whereby Rsu-1 containing the protein interacting LRR domain not only plays a significant role in the multi-protein assembly but also contributes to regulate a variety of signaling pathways and cellular processes during distinct spatiotemporal dynamics.

Given that the integrin-mediated adhesion complex comprises hierarchical clusters that control diverse sets of many biological processes, investigating the molecular basis of the interaction between Rsu-1 and PINCH-1 is crucial for understanding how the Rsu-1-mediated multi-protein assembly is orchestrated and what it can impact on the large architecture and cellular signaling. In the present study, we present the high-resolution crystal structure of Rsu-1 and its complex with a tandem repeat of LIM4-5 domains of PINCH-1. Together with biophysical and cellular experiments, we provide the molecular basis of target binding and specificity by Rsu-1. We also show that the interaction of Rsu-1 with PINCH-1 is essential for the localization to the sites of focal adhesion and the stabilization of the complex that is potentially implicated in the MAPK signaling pathway.

## Results

### Expression and Characterization of Rsu-1

To facilitate biochemical and structural studies, we established the protein expression and purification for the recombinant full-length Rsu-1 using baculovirus-insect cell system (Figure 1A). The biochemical characterization with size exclusion chromatography revealed a robust tight complex formation between the recombinant Rsu-1 and PINCH-1 LIM4-5 domains at 1:1 stoichiometry (Figure 1B-E), consistent with earlier report that the C-terminal LIM5 domain contains the binding site for Rsu-1 (8,12). We determined the binding affinity between Rsu-1 and PINCH-1 LIM4-5 domains at a single digit nanomolar (1.37±0.50 nM) using biolayer interferometry measurement (Figure 1F, Table 1). To gain structural insight into the architecture of Rsu-1, we first crystallized and solved its crystal structure by a single-wavelength anomalous dispersion (SAD) method using a single selenomethionine (SeMet)-substituted crystal. The structure of Rsu-1 was refined at 1.76-Å resolution to an R_factor_ of 19% and an R_free_ of 21% with good stereochemistry (Table 2).

**Table 1.** Summary of the binding interaction between Rsu-1 and PINCH domains.

| Binding interaction | Mean KD (nM) |
| --- | --- |
| Rsu-1 (WT):PINCH-1 LIM4-5 domain (WT) | $1.37 \pm 0.50$ |
| Rsu-1 (WT):PINCH-1 LIM 5 domain (WT) | $1.77 \pm 0.59$ |
| Rsu-1 (R2W):PINCH-1 LIM5 domain (WT) | $33,340 \pm 1,743$ |
| Rsu-1 (FRAA):PINCH-1 LIM5 domain (WT) | $1,386 \pm 100$ |
| Rsu-1 (WT):PINCH-2 LIM5S (WT) | $924.85 \pm 194.45$ |
| Rsu-1 (WT):PINCH-2 LIM5L (WT) | $5,952 \pm 560$ |
| Rsu-1 (WT):PINCH-1 LIM5 (D295V) | $1,520 \pm 200$ |

**Table 2.**
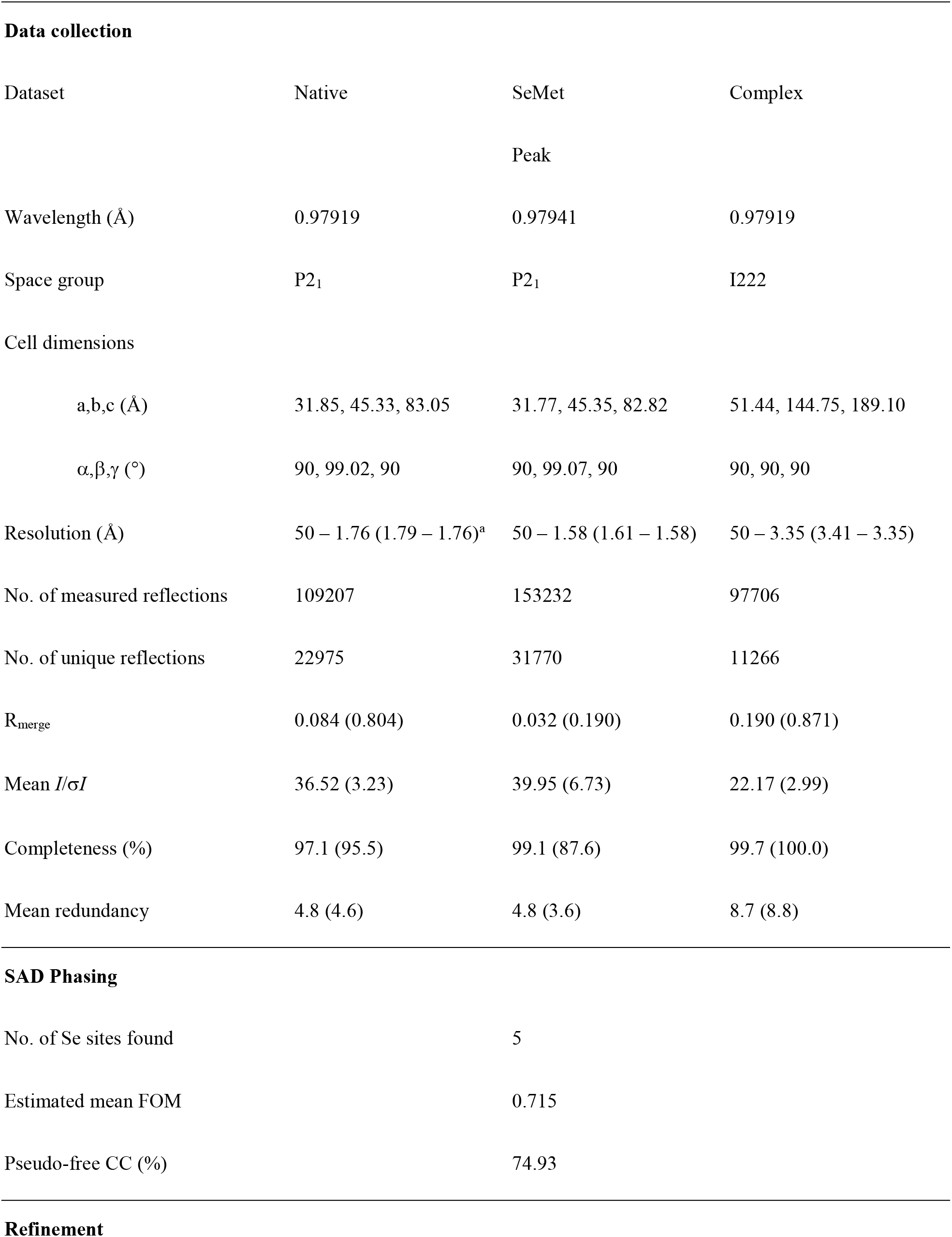
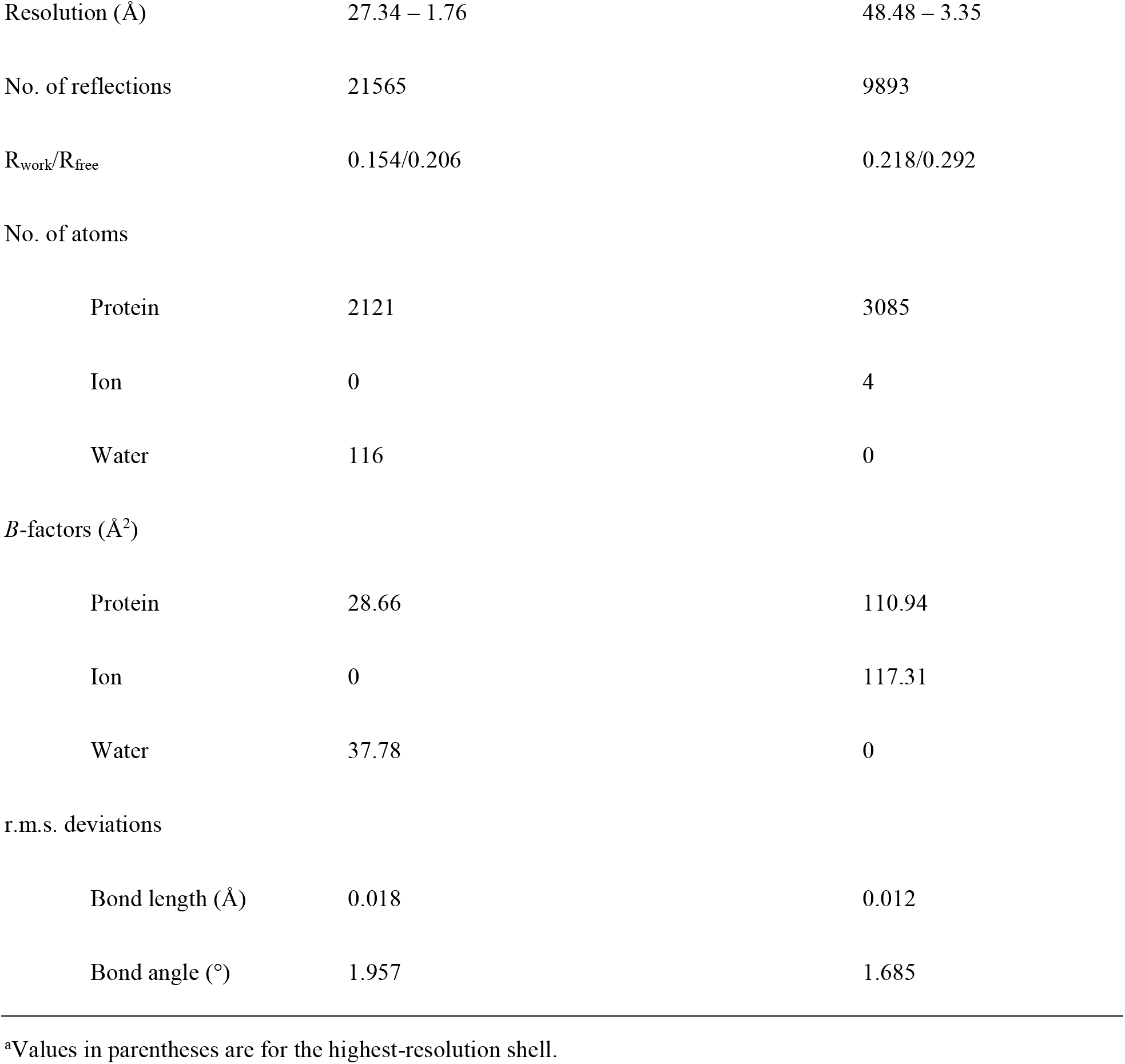
Data collection, phasing, and refinement statistics for Rsu-1 and its complex with PINCH-1 LIM4-5.

**Figure 1.**
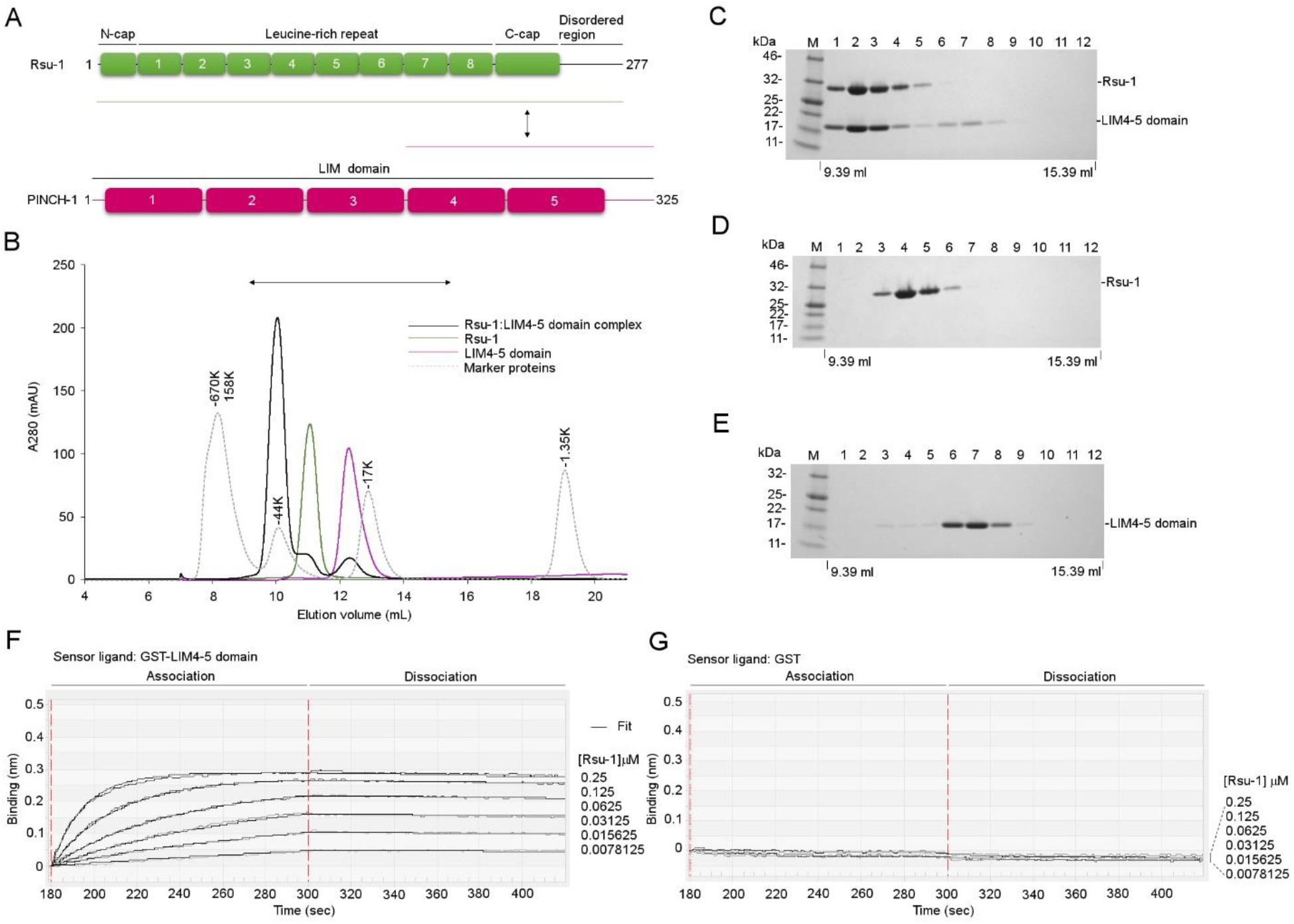
Characterization of the complex between Rsu-1 and PINCH-1. (A) Domain organization of Rsu-1 and PINCH-1. The construct regions used for crystallization experiments are denoted with green (Rsu-1) and magenta (PINCH-1) lines. (B) Gel filtration analysis of Rsu-1, PINCH-1 LIM4-5 domains, and its complex on size exclusion column with Superdex 75 Increase 10/300 GL. Four elution curves of Rsu-1 (green), PINCH-1 LIM4-5 domain (magenta), Rsu-1:LIM4-5 complex (black), and standard marker proteins (gray dot) are overlaid. The binary complex between Rsu-1 and PINCH-1 LIM4-5 domain elutes at a position similar to 44K marker protein. (C to E) Analysis of the eluents of twelve 500 μL fractions between 9.39 and 14.89 ml from an analytical Superdex 75 Increase chromatography column for Rsu-1, PINCH-1 LIM4-5 domain, and its complex. The SDS-PAGE gels were visualized with Coomassie staining. (F) Real-time measurement of the binding interaction between Rsu-1 and PINCH-1 LIM4-5 domain with BLItz system. GST-fused PINCH-1 LIM4-5 domain was immobilized onto anti-GST biosensor, and the binding of the indicated concentrations of Rsu-1 (mobile phase) was monitored in real-time. The association and dissociation phases are highlighted. (G) Control measurement with GST alone in the same concentrations of Rsu-1 as seen in (F).

### Overall structure of the leucine-rich repeat (LRR) protein Rsu-1

The structural analysis revealed that Rsu-1 adopts an arc-shaped solenoid architecture with the canonical LRR domain that comprises the N-terminal cap region followed by eight tandem LRRs and the C-terminal cap region with approximate dimensions of 80 Å × 40 Å × 30 Å (Figure 2A). The N-terminal cap region consists of one α-helix (residues from 5 to 15) and a loop, whereas the C-terminal cap region of Rsu-1 is comprised of the ninth β-strand followed by a helix-turn-helix motif. Amphipathic structural features in those N- and C-terminal cap regions shield the hydrophobic core residues in the first and last (eighth) LRRs to stabilize the structural integrity of the LRR domain, respectively. The terminal helix (residues 240 to 251) in the C-terminal cap region then folds toward the convex surface of the LRR region.

**Figure 2.**
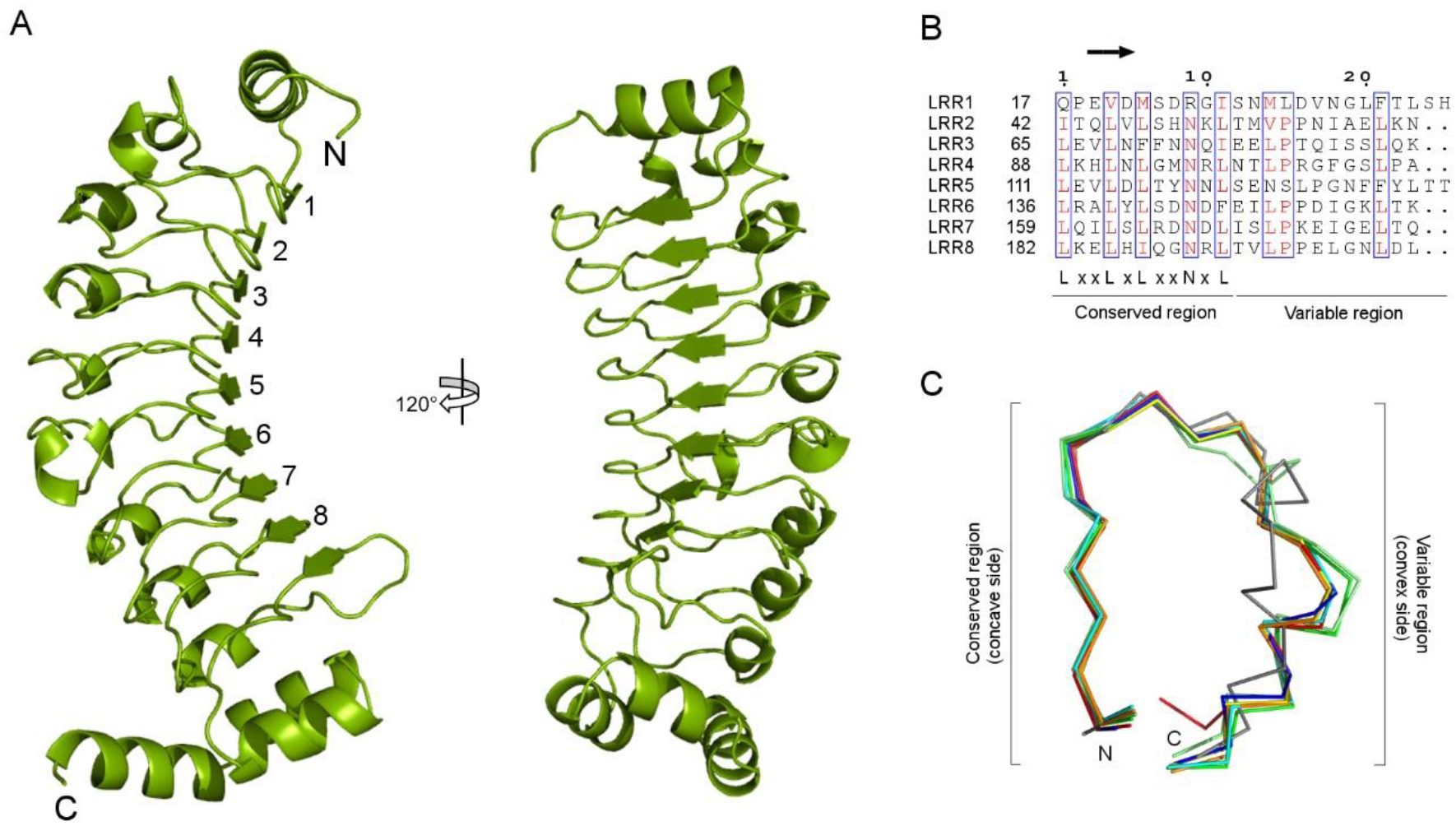
Crystal structure of the human Rsu-1. (A) Two orthogonal views of ribbon diagram of the crystal structure of Rsu-1. The structure of Rsu-1 includes residues from 1 to 252 but lacks the following C-terminal region (residues 253-277) owing to no visible electron density. (B) Alignment of individual LRR sequences of Rsu-1. Residues with high similarity are colored in red and framed in box. In Rsu-1, six LRR motifs (LRR2, LRR3, LRR4, LRR6, LRR7, and LRR8) contain one 3_10_ helix in each variable region, constituting to the 23-residue LRR sequence. By contrast, the first (LRR1) and fifth (LRR5) motifs contain two 3_10_ helices and two residue extension in their respective variable regions, resulting in the divergent motif with 25-residue LRR sequence. (C) Superposition of individual LRR fragments depicted in Cα trace models with distinct colors. Variable regions in LRR1 (gray) and LRR5 (light green) exhibit distinct conformations, whereas the conserved regions in all LRR fragments retain high structural regularity.

The LRR sequence of Rsu-1 consists of the consensus motif comprising LxxLxLxxNxLx_(*n*)_, where x denotes any residue and (*n*) represents the number of discrete residues, and the leucine residues are often substituted by other hydrophobic residues (10,11). The LRR sequence can be subdivided into two regions: conserved and variable regions (11). The conserved region in the LRR sequence of Rsu-1 consists of a short β-strand and a loop (Figure 2B). A parallel β-sheet with nine β-strands from eight LRR motifs and the following segment composes a characteristic concave surface in the LRR domain of Rsu-1 (Figure 2A). The variable region in the LRR motif of Rsu-1 is relatively divergent and features a distinctive convex surface (Figure 2B). Despite the substituted residues in the conserved region such as glutamine at the position 1, the LRR1 motif retains a secondary structure similar to other LRRs (Figure 2C), stabilizing the structural integrity of the LRR domain of Rsu-1. By contrast, the ninth β-strand in the following (eighth) LRR motif of Rsu-1 is aligned to a parallel β-sheet in the concave surface but a preceded four-residue insertion and a following helix-turn-helix module build up a distinct structural organization from the LRR motif, resulting in the structural part of the C-terminal cap that stabilizes the internal domain of LRR, as seen in other canonical LRR-containing proteins (11).

A structural database search with the DALI server (23) revealed a number of structures of LRR-containing proteins with divergent functions (Figure S1, Table S1). Apart from LRR-containing genes from human pathogen *Leptospira interrogans* such as LIC11098 that exhibits a high structural similarity with Z-scores of 26.6 (root-mean-square-deviation, RMSD, of 2.0 Å) (24), it is noteworthy that the structure of Rsu-1 resembles that of a human protein phosphatase 1 regulatory subunit 7 (a.k.a. SDS22) (25), a prototype of SDS22-like subfamily in the LRR superfamily (10) (Figure S1). Rsu-1 shares a 27% sequence identity with SDS22 that comprises 12 LRRs (25). The structure of Rsu-1 superimposes with that of SDS22 with an RMSD of 2.8 Å for the equivalent Cα atom pairs. Although Rsu-1 comprises two distinct variable sequence lengths (23- and 25-residues) in the LRRs that resemble in part those in the plant-specific (PS) subfamily in the LRR superfamily (10), our structural analysis strongly suggests that Rsu-1 falls into the SDS22-like subfamily rather than the PS subfamily (22). Other structurally related mammalian protein in the SDS22-like subfamily from the DALI search is the extracellular domain of mouse platelet receptor glycoprotein Ibα (GPIbα) (26). Those two representative SDS22 class LRR proteins bind their cognate partner proteins through their concave surfaces (27,28). Thus, it is of interest to understand how Rsu-1 with similar concave surface recognizes a different target (see below).

### Structure of the high affinity complex between Rsu-1 and PINCH-1 LIM4-5 domains

To understand the molecular basis of target recognition by Rsu-1, we set out to structurally characterize the complex between Rsu-1 and PINCH-1 LIM4-5 domains by X-ray crystallography. The high affinity complex between a tandem repeat of LIM4-5 domains of PINCH-1 and Rsu-1 was purified in multi-milligram quantities and subjected to crystallization screening. This high affinity complex enabled to grow crystals that diffracted at a moderate resolution of 3.35-Å, and the crystal structure of the complex was solved by molecular replacement (Table 2).

The structural analysis revealed that the PINCH-1 LIM4-5 fragment binds via its LIM5 domain to the concave surface of a parallel β-sheet in the LRR domain of Rsu-1 (Figure 3A). The pattern of the binding mode resembles those in other LRR-protein ligand bound structures such as a complex between follicle-stimulating hormone and its receptor with LRR domain (29) and a complex between NetrinG and NetrinG ligand 1 with LRR domain (30), referring to as “hand-clasp”. The superposition of the structure of Rsu-1 bound form with that of its unbound form showed a root-mean-square deviation (r.m.s.d.) of ∼0.6-Å for the Cα atom pairs, indicating that Rsu-1 retains its rigid architecture and does not undergo major structural change upon the binding to PINCH-1. The conformation in the C-terminal region of Rsu-1, which is disordered in the unbound form, essentially remains the same structural arrangement as the unbound form, suggesting that the C-terminal region is not involved in the direct binding to PINCH-1. The binding interface is relatively extensive and continuous, and buries approximately 2,400 Å_2_ of solvent-accessible surface area (Figure 3B-E) that is comparable to those found in the LRR-protein ligand complexes at high affinity (11). The goodness of fit in the interface characterized by a shape correlation statistic (Sc) (31) reveals that the Sc value is measured at a shape complementarity of 0.62 at the interface of the Rsu-1-LIM4-5 complex. The fitness by shape complementarity itself may be somewhat moderate when compared to antibody-protein complex and enzyme complex (31) but a relatively large buried surface through extensive concave surface in the LRR domain may compensate to accommodate protein ligand, resulting in high affinity binding (11). The mechanism for the interaction in the Rsu-1-LIM complex resembles those seen in other LRR-protein ligand complexes such as the complex between platelet receptor GPIbα and vWF-A1 (Sc, 0.60) (27), the complex between follicle-stimulating hormone and its receptor (Sc, 0.59) (29), the complex between NetrinG and NetrinG ligand 1 (Sc, 0.58) (30), and the complex between SDS22 and protein phosphatase 1α catalytic domain (Sc, 0.57) (28) (Figure S2). Those LRR-protein ligand complexes display relatively large buried surfaces despite moderate shape complementarity measurements, and the strength of the interaction for each complex is maintained by specific interactions such as electrostatic interactions and hydrogen bonds (see below) (11).

**Figure 3.**
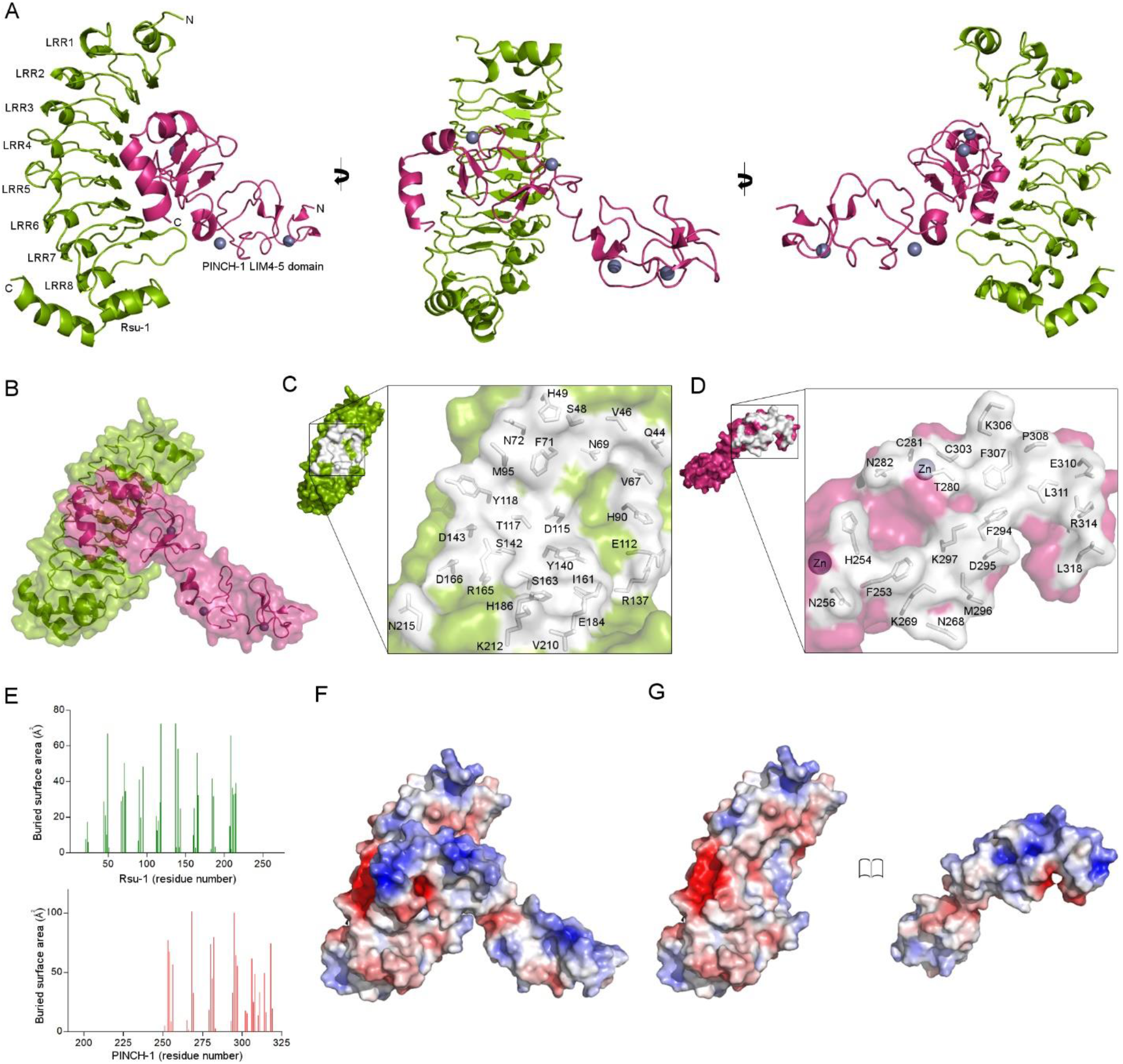
Crystal structure of the complex between Rsu-1 and PINCH-1 LIM4-5 domain. (A) Three orthogonal views of ribbon drawing of the crystal structure of the complex between Rsu-1 (green) and PINCH-1 LIM4-5 domain (magenta). Zinc atoms are depicted in spheres. (B) Transparent surface and cartoon model of the crystal structure of the complex. (C-D) Open molecular surface views on the binding interfaces between Rsu-1 (green) and PINCH-1 LIM4-5 domain (magenta). The interfacial residues are depicted in stick models. (E) Analysis of the buried surface area in the binding interface between Rsu-1 (green) and PINCH-1 LIM4-5 domain (red). (F) Electrostatic surface potential map of the complex at the same view of (B). (G) Open views of the electrostatic surface potential maps of Rsu-1 (left) and PINCH-1 LIM4-5 domain (right).

### The binding mode for the high affinity complex of Rsu-1 with the LIM4-5 domains

The analysis of electrostatic surface potential of the Rsu-1-LIM complex reveals that each protein surface displays distinct complementary charge distributions and binding architecture at the interface (Figure 3F-G). Two polar residues of Arg165 and Asp166 from the LRR β7 strand of Rsu-1 that are conserved across species play a prominent role in the interface, by forming salt-bridges with Asp295 and Arg314 of PINCH-1 LIM5 domain, respectively (Figure 4). Strikingly, the identification of the salt-bridge between Arg165 of Rsu-1 and Asp295 of PINCH-1 is complementary and in agreement with the previous genetic study in *Drosophila*, which showed that the residue (Asp303) of fly PINCH-1 corresponding to Asp295 in the human orthologue was crucial for the binding to Rsu-1 (32). A number of charged residues also provide additional salt-bridges and hydrogen bonds in the interface (Figure 4B). Beside the hydrogen bonds and salt-bridges, two hydrophobic residues of Phe71 and Met95 of Rsu-1 that contact to Phe307 and Pro308 of PINCH-1 LIM5, respectively, also significantly contribute to the binding interaction (Figure 4C). Taken together, those highly specific interactions allow PINCH-1 to dock onto a relatively rigid concave surface of Rsu-1, and define the conserved binding mode and architecture of the complex (Figure 4D-E).

**Figure 4.**
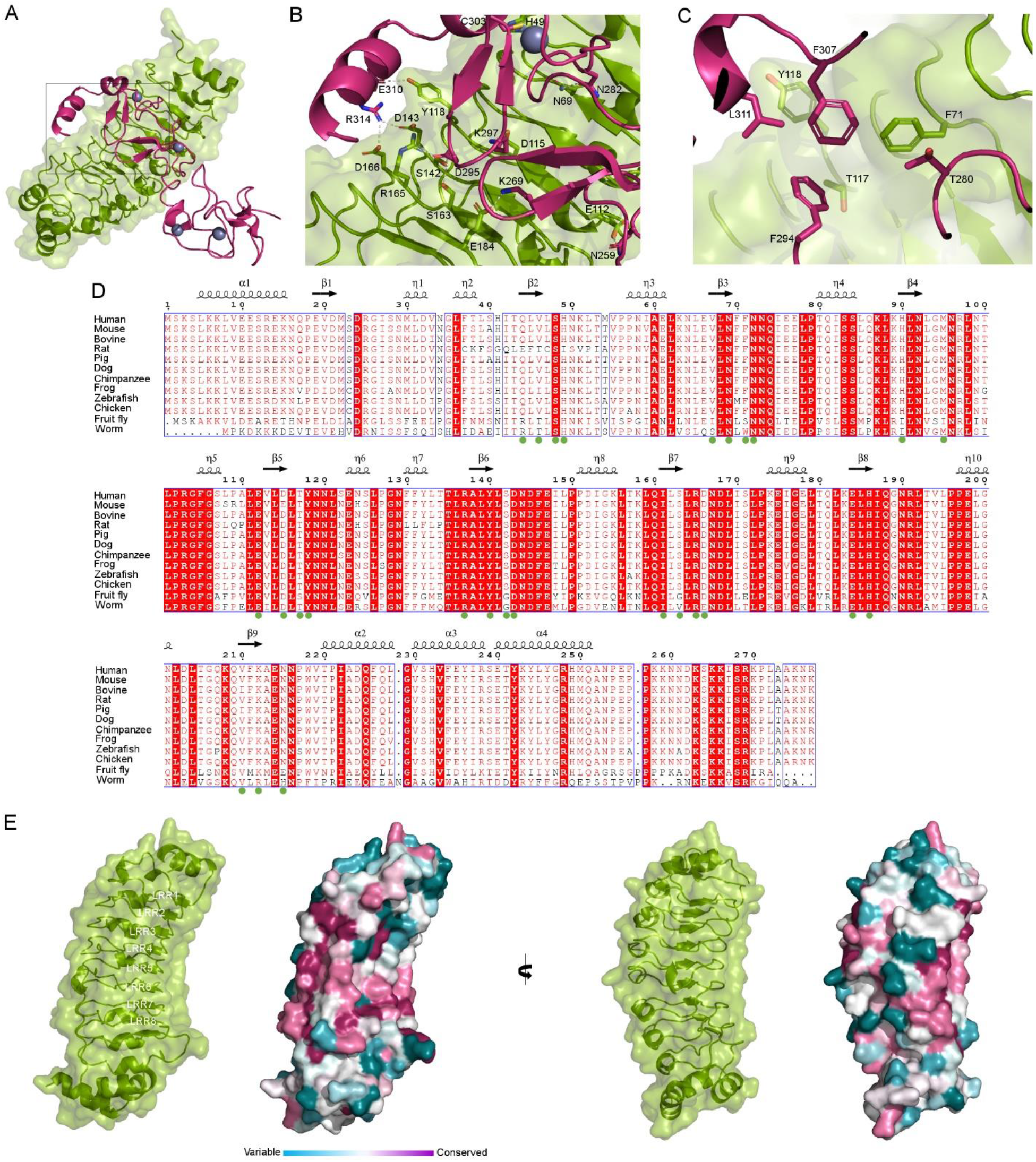
Determinants of the specific binding interaction and conservation analysis for the complex between Rsu-1 and PINCH-1 LIM4-5 domain. (A) Overall structure of the complex depicted in transparent surface and ribbon models. Rsu-1 and PINCH-1 LIM4-5 domain are colored in green and magenta, respectively. The close-up view area is framed. (B) A close-up view of the selected contact residues for inter-molecular side-chain hydrogen-bonding and salt-bridge interactions in the interface. Zinc atom is depicted in spheres. (C) A close-up view of selected contact residues for inter-molecular hydrophobic interactions in the interface. (D) Multiple sequence alignment of Rsu-1 across species with the Clustal Omega program followed by rendering with the program ESPript. The residues for the PINCH-1-binding are depicted in filled green circles below the sequence. The secondary structures are depicted above the sequence. η represents 3_10_ helix. The invariant residues are highlighted as white characters on red background, whereas the highly conserved residues are shown in red characters. (E) Surface representation of Rsu-1 colored by sequence conservation across species at two orthogonal views. A transparent surface model, along with ribbon drawing of Rsu-1, is also depicted.

To identify the functionally important specific binding residues involved in the interface, a series of alanine substitutions in both Rsu-1 and the PINCH-1 LIM5 domain were generated based on the algorithm of the Protein Interfaces, Surfaces and Assemblies (PISA) (33). The recombinant mutant proteins of the PINCH-1 LIM5 domain were expressed and evaluated for their interaction with Rsu-1 using *in vitro* binding experiments, revealing impaired binding capability for the following mutants: Asp295Ala, Phe307Ala, and Arg314Ala. Using biolayer interferometry assessment, we next measured the binding kinetics and affinity between Rsu-1 and PINCH-1 LIM5 and confirmed that Rsu-1 indeed binds the PINCH-1 LIM5 domain at a single-digit nanomolar affinity (Figure 5, Table 1) that is comparable to the binding affinity with PINCH-1 LIM4-5 domains (Figure 1F, Table 1). The control experiments indicated that Rsu-1 at the concentration range from 1 to 16 μM did not bind GST-loaded biosensors (Figure 5D), demonstrating that the slow off rate was not derived from non-specific binding to the sensors. This high affinity complex is in great agreement with size exclusion chromatography experiments that allow to co-purify as a 1:1 stoichiometric complex (Figure 5G). Those results are consistent with the structural results that the PINCH-1 LIM5 domain solely engages Rsu-1 with considerable binding interactions through charge and shape complementarity. Double alanine substitutions of Rsu-1 at Phe71 and Arg165 (designated as FRAA) resulted in a significantly reduced binding to the PINCH-1 LIM5 at over 1,000-fold impaired affinity (∼ 1 μM) (Figure 5B, Table 1). To further validate the binding function, the side chains of Phe71, Arg165, and Asp166 of Rsu-1 were substituted by arginine and tryptophan with larger and counter nature. We found that the triple mutant (Phe71Arg, Arg165Trp, and Asp166Arg; designated as R2W) of Rsu-1 dramatically impaired the binding interaction to the PINCH-1 LIM5 domain, confirming the central role of those hydrophilic and hydrophobic residues in the interface (Figure 5C, Table 1). Correct protein folding of the triple mutant (R2W) of Rsu-1 was assessed by retained high levels of protein expression and solubility with monodisperse in solution (Figure 5I). As a reciprocal binding experiment, the conserved Asp295 in the PINCH-1 LIM5 domain that makes a salt bridge with Arg165 in Rsu-1 was substituted by Val295 according to the previous genetic study (32). Using the BLItz binding experiment, we found that the single residue substitution in the PINCH-1 LIM5 domain reduced the binding interaction to Rsu-1 (Table 1). Taken together, those results strongly suggest that those two distinct regions comprising hydrophobic contact, along with hydrogen bonds and salt-bridges, play a major role in the structural determinant of the interaction between Rsu-1 and PINCH-1.

**Figure 5.**
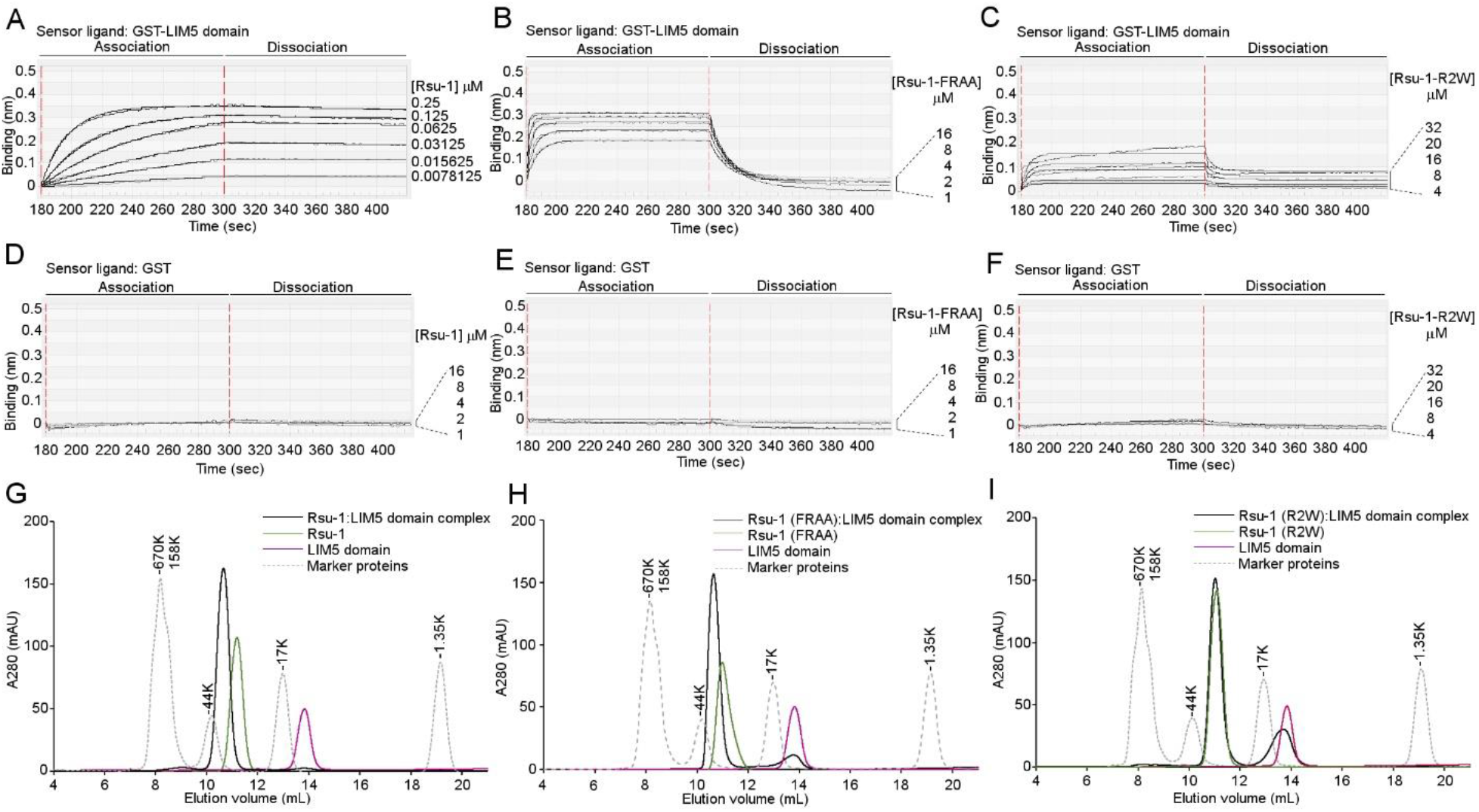
The binding interaction analysis of the Rsu-1 mutant proteins with PINCH-1 LIM5 domain. (A) Real-time binding analysis of the interaction between Rsu-1 (WT) and PINCH-1 LIM5 domain with biolayer interferometry (BLItz) measurement. Representative binding curves at dose-dependent concentration of analyte (Rsu-1) are shown. (B) The BLItz binding data of the interaction between the Rsu-1 double mutant protein (FRAA) and PINCH-1 LIM5 domain. (C) The BLItz binding data of the interaction between the Rsu-1 triple mutant protein (R2W) and PINCH-1 LIM5 domain. (D) Control experiment of the BLItz measurement with GST alone on the sensor chips. (E) Overlaid elution profiles of the complex (Rsu-1 WT:PINCH-1 LIM5 domain) and each unbound protein (Rsu-1 WT or PINCH-1 LIM5 domain) with analytical size exclusion chromatography on a Superdex 75 Increase 10/300 column, calibrated with standard marker proteins. The complex was eluted at 1:1 stoichiometric complex. (F) Overlaid elution profiles of the loss-of-binding triple mutant (R2W) of Rsu-1 and PINCH-1 LIM5 domain on the Superdex 75 Increase column. The triple mutant protein (R2W) did not form a stable complex with PINCH-1 LIM5 domain at 1:1 molar ratio, confirming the determinant for the major contact residues.

### Structural insight into isoform-specific interaction of PINCH with Rsu-1

PINCH comprises two isoforms in human (34). Both two isoforms (PINCH-1 and -2) share a high degree of protein sequence identity (∼86%) with similar domain architecture consisting of a tandem repeat of five LIM domains except for the C-terminal LIM5 domain that has the 11-residue extension in PINCH-2 than in PINCH-1. Beside the C-terminal extension, the sequences of the PINCH-2 LIM5 domains are highly conserved across species (Figure 6A). Strikingly, 18 out of 20 direct contact residues of human PINCH-1 LIM5 domain to Rsu-1 are invariant to the human PINCH-2 LIM5 domain (Figure 6A). Two variable residues of Tyr258 and Asn259 in the LIM5 domain of PINCH-2 are relatively comparable in size and charge to Phe253 and His254 in that of PINCH-1 (Fig. 6A). This argues with a previous study that PINCH-1 binds Rsu-1 but PINCH-2 does not (12). To investigate a structural basis of isoform specific interaction by Rsu-1, the recombinant PINCH-2 LIM5 protein with the C-terminal 11-residue extension (designated as PINCH-2 LIM5L) that involves the region equivalent to the PINCH-1 LIM5 domain was bacterially expressed, and the binding interaction with Rsu-1 was examined by biolayer interferometry assay. We found that while the PINCH-2 LIM5L interacted with Rsu-1, the affinity is at low micromolar affinity of 5.95 μM (Figure 6B) that is more than 3,000 times weaker than that of PINCH-1 LIM5 domain (Figure 5A). Given highly conserved sequences among two PINCH proteins across species, the overall architecture and structure elements of the core region of those LIM5 domains would be expected to be highly homologous. On the other hand, the C-terminal 11-residue extension of the PINCH-2 LIM5 domain differs from that of PINCH-1 (Fig. 6A), which made us wonder whether this region affects the PINCH-2 binding to Rsu-1. We then expressed the shorter fragment of PINCH-2 LIM5 short domain without the C-terminal 11 residues (designated as PINCH-2 LIM5S) and evaluated the binding interaction with BLItz system. We found that the PINCH-2 LIM5S bound to Rsu-1 at approximately 6-times higher affinity than PINCH-2 LIM5L (Figure 6C). The purified PINCH-2 LIM5S protein exhibited a soluble monodisperse condition in solution and surprisingly formed a robust complex with Rsu-1 at 1:1 molar ratio (Figure 6D). Those results suggest that an isoform-specific C-terminal 11-residue extension of PINCH-2 seems to determine the binding selectivity to Rsu-1 among PINCH proteins despite the highly conserved residues in the LIM5 domains between PINCH-1 and PINCH-2 (Figure 6A). Molecular modeling analysis further confirmed that the interaction between the PINCH-2 and Rsu-1 depends on the highly conserved residues in the LIM5 domain, and that Asp300 of the human PINCH-2 equivalent to Asp295 of the human PINCH-1 may make a significant interaction of salt-bridge formation with Arg165 of Rsu-1. Thus, the unexpected potent binding of PINCH-2 LIM5S to Rsu-1 is likely mediated by the same conserved residues in PINCH-1 LIM5. As our modeling analysis and secondary structure prediction did not reveal the structural information and conformation of the C-terminal extension of PINCH-2 LIM5 domain, what the C-terminal extension of PINCH-2 looks like and how it is far from the contact residues to Rsu-1 for steric hindrance could be interesting questions. On the other hand, the C-terminal helix in the LIM5 is relatively in close proximity to the C-terminal cap in the LRR domain of Rsu-1 upon association. Thus, it is expected that the 11-residue extension in the PINCH-2 LIM5 domain may generate a potential steric hindrance upon association with Rsu-1, which provides a basis for understanding the selectivity for the binding to Rsu-1 between PINCH-1 and PINCH-2. Despite those biochemical and structural findings, it should be noted that the interaction of Rsu-1 with PINCH-2 is extremely weaker than those with PINCH-1, and its functional significance in the biological context as a counter pair at the sites of focal adhesion remains unknown.

**Figure 6.**
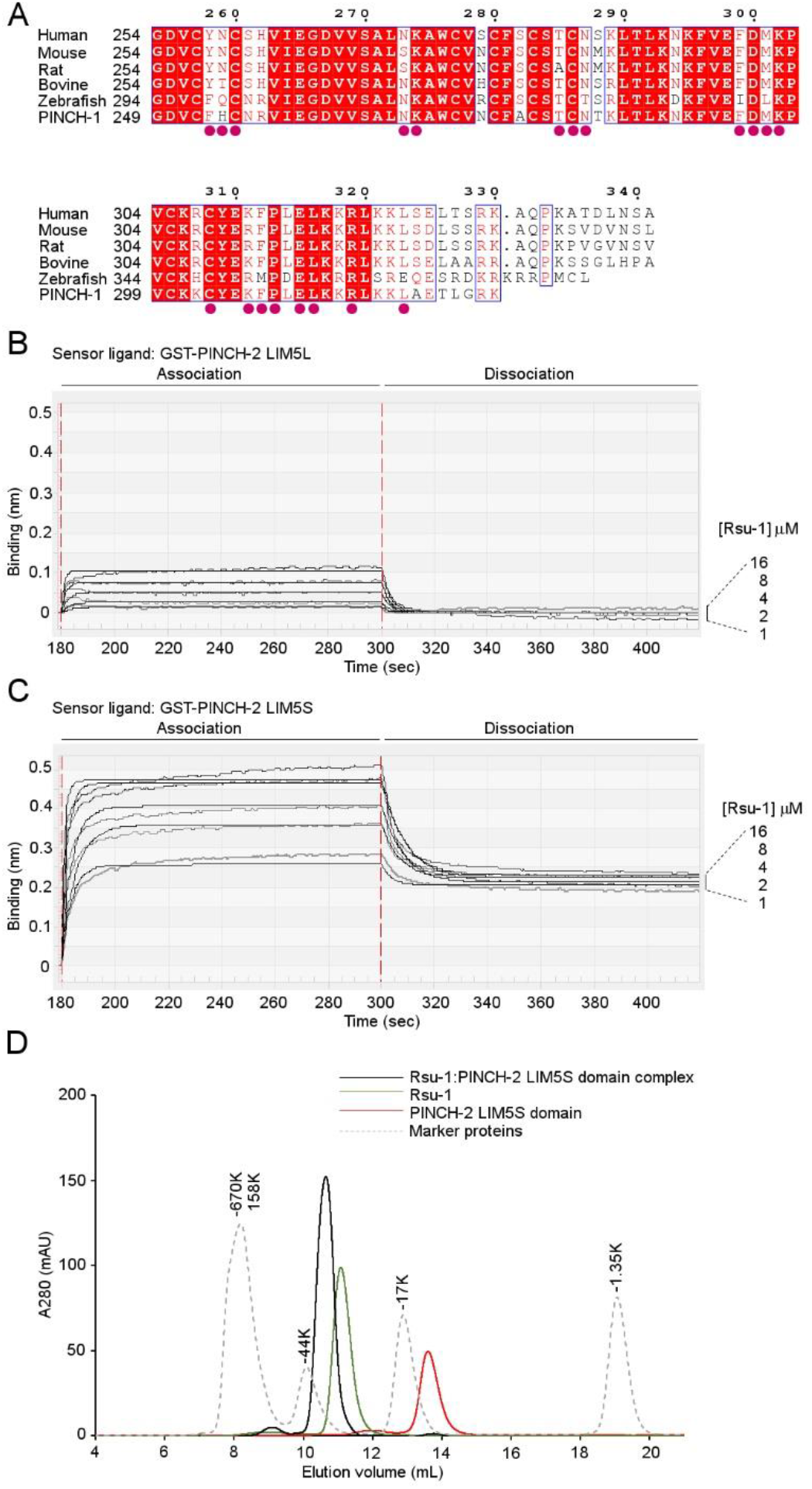
Biochemical analysis of the isoform specific interaction between Rsu-1 and PINCH-2. (A) Multiple sequence alignment of PINCH-2 across selected vertebrate species and comparison with the human PINCH-1 sequence. The numbering above the alignment refers to the human PINCH-2 sequence. The invariant residues are highlighted in white on red background, whereas the highly conserved residues are shown in red characters and framed. The contact residues of the human PINCH-1 LIM5 domain to Rsu-1 are depicted in filled red circles. (B) The BLItz sensorgrams for the interaction between Rsu-1 and PINCH-2 LIM5L. (C) The BLItz sensorgrams for the interaction between Rsu-1 and PINCH-2 LIM5S. The shorter fragment of PINCH-2 LIM5S without the C-terminal 11-rsidue extension exhibited higher binding affinity than larger fragment (PINCH-2 LIM5L). (D) An overlaid elution profile of the PINCH-2 LIM5S domain, Rsu-1 wild type, and their complex by size exclusion chromatography on Superdex 75 Increase column.

### Functional implications for the interaction between Rsu-1 and PINCH-1

Having demonstrated the robust high affinity complex and isoform-specific interaction between Rsu-1 and PINCH-1, we next sought to investigate functional significance of this interaction. A series of constructs of Rsu-1 to various fusion tags such as GFP, Myc, and HA were generated and transiently expressed in model cell lines including HeLa, MCF10A, and MEF, and their abilities to localize at the focal adhesion sites were examined using confocal microscopy. Our confocal microscopy experiments showed that when expressed as a GFP fusion protein in HeLa cells Rsu-1 appeared in mostly cytosolic fraction and occasionally in close proximity to nucleus but not at the sites of focal adhesion despite its robust high affinity complex formation with PINCH-1. Although a nuclear localization signal is not found in Rsu-1, there might be an additional as-yet-uncharacterized function of Rsu-1 that shuttles through its associated proteins to nucleus, which resembles other LRR-containing proteins such as ERBIN (35) and a nuclear protein SDS-22 (36). Since Rsu-1 tightly binds to PINCH-1, exogenously expressed RSU-1 may not effectively replace the endogenous Rsu-1 to bind PINCH-1. We then knocked down Rsu-1 in MCF10A cells and re-expressed Myc-tagged Rsu-1 in the cells. The Myc-tagged wild type Rsu-1 localized to the focal adhesion sites but the triple mutant (R2W) of Rsu-1 that abrogates the binding to PINCH-1 did not(Figure 7A). Our results are consistent with previous studies that pEGFP-Rsu-1 and Myc-tagged PINCH-1 co-localized at the sites of focal adhesion in Cos-7 cells (12). Those results demonstrate that the localization of Rsu-1 at the sites of focal adhesion in mammalian culture cells is significantly mediated by its LRR domain that binds the C-terminal LIM5 domain of PINCH-1, and that constitutively expressed endogenous protein can interfere with the functionality of its exogenously expressed fusion-tagged protein when the binding affinity is extremely high (at a single digit nanomolar affinity).

**Figure 7.**
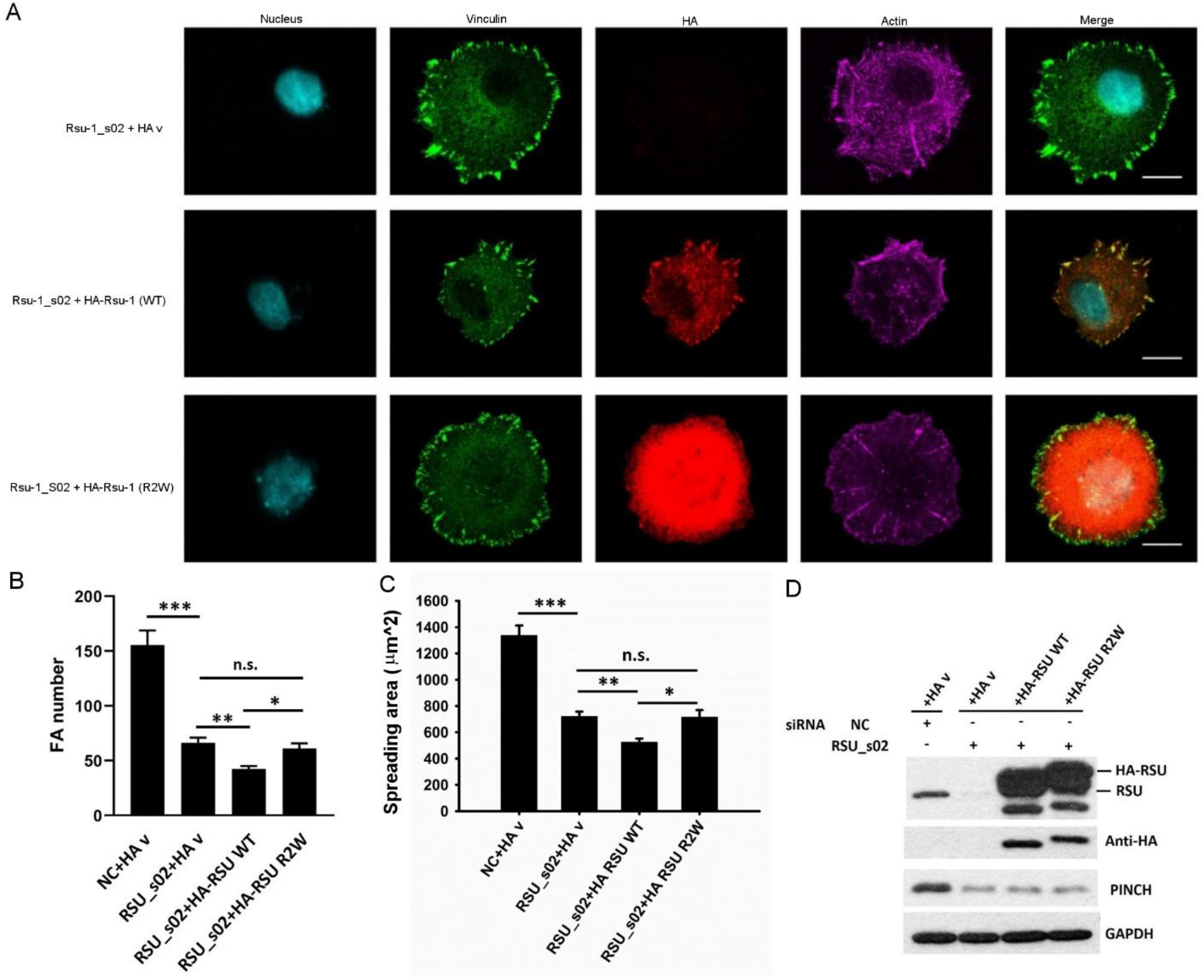
Functional significance of the interaction between Rsu-1 and PINCH-1 at the focal adhesion sites. (A) Co-localization of HA-tagged Rsu-1 wile type (WT) or triple negative mutant (R2W) at the vinculin-containing focal adhesion sites in siRNA treated MCF10A cells. HA-tagged Rsu-1 WT was co-localized to the vinculin-containing focal adhesion sites, whereas the triple mutant (R2W) was not. Transfected cells were allowed to spread on fibronectin coated coverslips for 2 hours before staining. Images were captured with confocal microscope, under 63x magnification. Scale bar, 10 µm. (B) Quantitative analysis of the focal adhesion number in MCF10A cells, showing that Rsu-1 WT resulted in less focal adhesion number than the triple mutant (R2W). (***P<0.001, **P<0.01, *p<0.05, N=60) (C) Quantitative analysis of spreading of MCF10A cells, showing that Rsu-1 regulated cell spreading through PINCH-1 binding. (***P<0.001, **P<0.01, *p<0.05, N=50). All values are given as mean ± S.E.M.

We next evaluated whether the interaction between Rsu-1 and PINCH-1 can impact on the formation of focal adhesion and cell spreading since previous study showed that Rsu-1 is essential for focal adhesion assembly and cell spreading (37). MCF10A cells under Rsu-1 siRNA treatment were transfected with either the HA-tagged wild type or a PINCH-1-binding deficient triple mutant (R2W) Rsu-1. Interestingly, the WT Rsu1-expressing cells in Rsu-1-depleted MCF10A had significantly less FAs than the Rsu1 R2W-expressing cells (Figure 7B). Correspondingly, the WT Rsu1-expressing cells exhibited significantly less spreading than the Rsu1 R2W-expressing cells (Figure 7C). It is noticeable that the protein level of endogenous PINCH-1 was reduced when MCF10A cells were treated with Rsu-1 siRNA, and that the reduction was not restored upon transiently expressing the wild type Rsu-1 (Figure 7D). Thus, it is conceivable that the reduction of focal adhesion number followed by cell spreading could coincide with the down regulation of endogenous protein of PINCH-1, suggesting a mutual relationship between transcriptional regulation and association through the Rsu-1-PINCH-1 axis. To circumvent those issues, we knocked down Rsu-1 in a PINCH-1-deficient HeLa cells, as previously demonstrated (38). Either PINCH-1/wild type Rsu-1 or PINCH-1/double negative mutant (FRAA) Rsu-1 was then co-expressed in those cell lines. Consistent with Figure 7B, cell spreading in the absence of PINCH-1 protein was also significantly reduced when the wild type Rsu-1 expressing cells were compared with the double mutant Rsu-1 expressing cells (Figure S3A-B). These data were surprising, but suggest that proper Rsu1/PINCH-1 levels may be critical for fine-tuning FA assembly process and FA-dependent cell adhesive processes such as cell spreading. Overall, the data demonstrate that Rsu-1 binding to PINCH-1 is crucial for the recruitment of RSsu-1 to focal adhesion sites to regulate cell adhesion dynamics.

### Integration of Rsu-1 within consensus adhesome machinery and implications for its potential scaffolding function

To provide mechanistic insights into the Rsu-1-mediated focal adhesion assembly, we next investigated the conformational stability of the interaction between Rsu-1 and the PINCH-1 LIM5 domain by protein thermal-shift assay. Our data revealed that the calculated melting temperature (T_m_) of Rsu-1 bound to PINCH-1 LIM5 domain (Tm=67.7±0.5°C) was significantly higher (11.6°C) than that in the absence of the protein ligand (Tm=56.1±0.2°C), suggesting a significant preference of the interaction for the increased protein stability (Figure 8A). As PINCH-1 is one component out of the evolutionarily conserved IPP heterotrimer adhesion machinery (39), it is intriguing whether Rsu-1 can be integrated in the IPP-containing adhesion complex. We next recombinantly expressed and purified the IPP heterotrimer complex (38) and kindlin-2 (40) that are involved in a consensus integrin adhesome (5), and analyzed *in vitro* reconstituted multi-protein complex formation by size exclusion chromatography. Our biochemical characterization has revealed that ILK, PINCH-1, α-Parvin, Kindlin-2, and Rsu-1 assembled into a heteropentamer protein complex (KIPPR) (Figure 8B-C). In this heteropentamer complex, the C-terminal LIM5 domain of PINCH-1 provides the binding site for Rsu-1, whereas the N-terminal LIM1 domain binds the N-terminal ankyrin repeat domain of the pseudokinase protein ILK (41). The C-terminal pseudokinase domain of ILK then concomitantly bridges α-Parvin (42) and Kindlin-2 (40), playing the central role in the hub of the KIPPR complex formation. (Figure 8D). Those results strongly suggest that Rsu-1 significantly contributes to the stabilization of the KIPPR complex formation that is supported by previous genetic results (20), and the KIPPR complex plays a crucial role in the integrin-mediated cell adhesion network.

**Figure 8.**
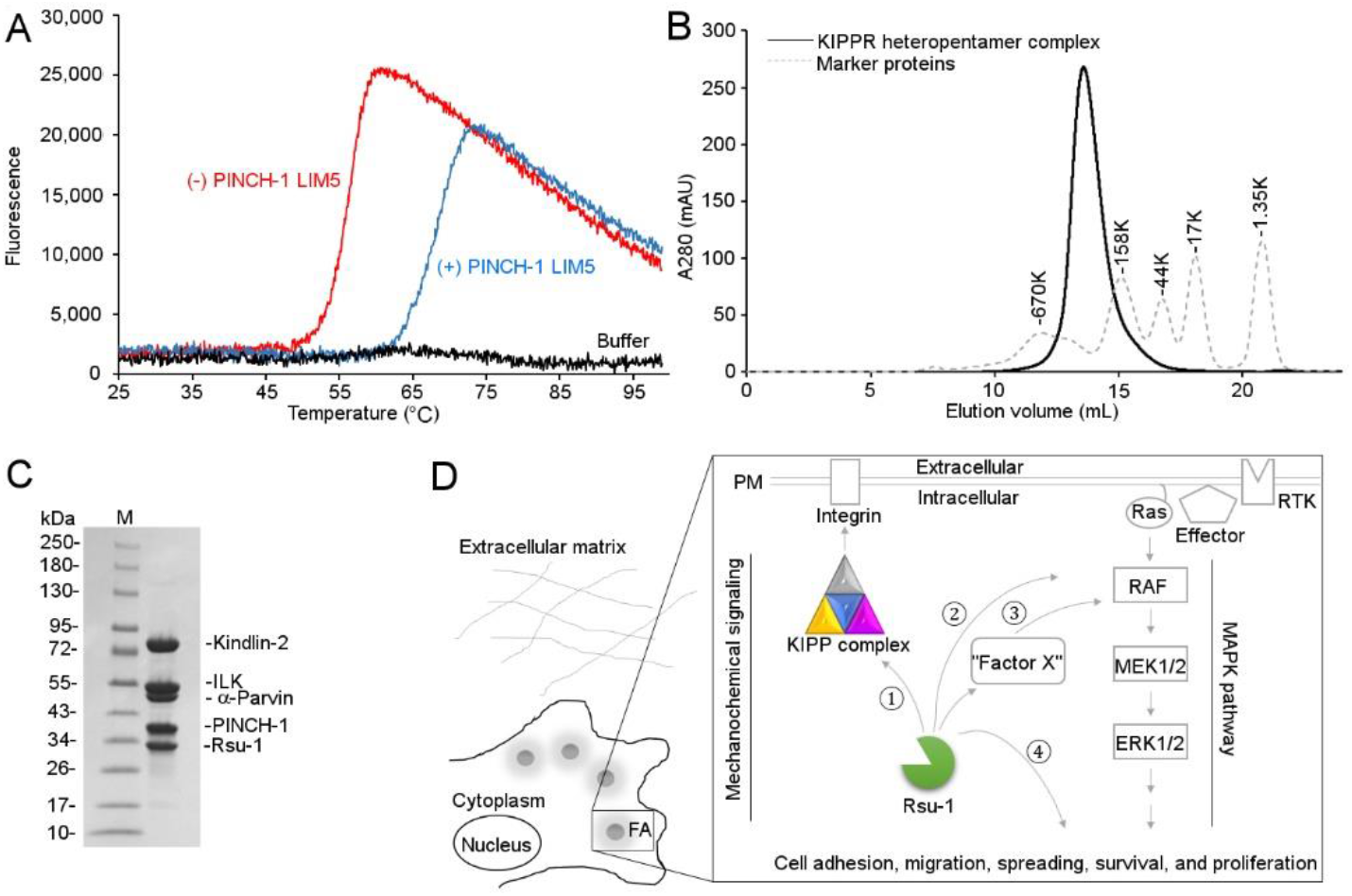
Impact of the interaction between Rsu-1 and PINCH-1 on the stabilization and maturation of the focal adhesion assembly. (A) Representative thermal denaturation profile of recombinantly purified Rsu-1 in the absence (red) and presence (blue) of the PINCH-1 LIM5 domain. The thermal profile of buffer with a fluorescent dye (as a no protein control) is overlaid. The thermal profile of PINCH-1 LIM5 is omitted due to flat low fluorescence signals. (B) An elution profile for recombinantly expressed and reconstituted Rsu-1-mediated heteropentamer protein complex using size exclusion chromatography on a Superose 6 column. The IPP heterotrimer complex comprising ILK, α-parvin, and PINCH-1 stably binds kindlin-2, forming a heterotetramer complex (KIPP). Addition of Rsu-1 to this heterotetramer complex (KIPP) results in a stable heteropentamer complex (KIPPR). The elution curve of the standard marker proteins is overlaid. (C) Coomassie-stained SDS-gels of the KIPPR heteropentamer complex from the major peak fraction eluted at 13.6 ml on a Superose 6 column. (D) Interaction network of Rsu-1 and its related molecules within consensus adhesome complex. A conceptual framework of Rsu-1-mediated interaction and potential regulation is illustrated in distinct pathways. Rsu-1 interacts with PINCH-1 at high affinity for the conformational stability and appears to constitutively bind the KIPP complex, resulting in the formation and stabilization of the heteropentamer complex for mechanochemical signaling (➀). Rsu-1 was previously claimed to interact with Raf-1/CRAF (➁) but the interaction may be by an indirect manner that may require as-yet-uncharacterized interacting molecules (“Factor X”) (➂). Rsu-1 may also potentially regulate the downstream event in the MAPK pathway linking to various cellular functions (➃).

As Rsu-1 was originally identified as a suppressor for the Ras-transformed cells (7), we also wondered whether Rsu-1 may function as a scaffolding protein to coordinate signaling such as the Ras-MAPK (Raf-MEK-ERK) signal transduction pathway that regulates cell growth, proliferation, differentiation, and apoptosis (43). Since Raf-1 (CRAF) is the first effector for the downstream of Ras linking to the MAPK pathway (44), it is intriguing whether Rsu-1 is involved in the Ras-Raf-1 axis. Noteworthy is that Raf-1 was previously shown to interact with Rsu-1 by *in vitro* binding experiment (19). Thus, it is conceivable that Raf-1 can potentially physically link Rsu-1 and Ras proteins, thereby the resultant signaling complex may regulate the Ras-mediated MAPK signaling pathways (45). We first explored to quantitate a potential interaction between Rsu-1 and Raf-1 using BLItz binding system but our initial efforts to generate recombinant full-length Raf-1 using bacterial and baculovirus expression systems failed owing to heterogeneous solubility issues. By contrast, a minimal fragment (CR1) comprising RBD and CRD of Raf-1 was successfully generated using bacterial expression system and utilized to examine a potential interaction with Rsu-1. Our quantitative assessment with BLItz biosensor system revealed that while Raf-1 CR1 domain significantly bound HRAS (G12V) that is consistent with recent binding study (46), it did not interact with Rsu-1 at a physiologically meaningful affinity (data not shown), suggesting that the CR1 does not involve the binding site for Rsu-1. Confirmation of a potential physical interaction between Raf-1 and Rsu-1 and the presence of additional as-yet-uncharacterized linker molecules should await further investigation until a full-length Raf-1 protein and a diverse set of interactors become available.

## Discussion

The structural and functional studies presented in this work provide the atomic view of a previously understudied focal adhesion protein Rsu-1 from the LRR superfamily involved in the integrin-mediated adhesion complex and reveal a novel architecture of the LRR-LIM protein complex. Our structural results reveal that Rsu-1 folds into a rigid arc-shaped architecture with eight consecutive LRRs shielded by two terminal capping modules and it engages via its extensive conserved concave surface the LIM5 domain of PINCH-1 in a pattern analogous to some LRR-protein ligand complex formation that is referred to as “hand-clasp” (29,30). We identify the binding interface of the high affinity complex between Rsu-1 and PINCH-1. Highly conserved salt-bridges, hydrogen bonds, and hydrophobic interactions are essential components for the specificity of the interaction and the formation of the high affinity complex between Rsu-1 and PINCH-1. Specifically, Arg165 in the LRR domain of Rsu-1 forms a key salt-bridge to Asp295 in the LIM5 domain of PINCH-1. These specific residues for the salt-bridge interaction are highly conserved across species, and the significance of the salt-bridge interaction is in agreement with a genetic study that revealed a critical role in the muscle hypercontraction in *Drosophila* (32). Interestingly, although a cluster of conserved interfacial residues of PINCH-1 for the binding to Rsu-1 is highly conserved in another isoform PINCH-2, the C-terminal region of the PINCH-2 LIM5 domain exhibits a significantly extended sequence, which disfavors the binding to Rsu-1 as shown in this study that is consistent with the results in mammalian culture cells (12). In this context, it is noticeable that some PINCH-1 proteins from some species also exhibit similar extended sequences in the C-terminal region (Figure S4). Based on our structural analysis, we propose that such extended region in the C-terminus of PINCH-2 LIM5 sterically restrains an inter-molecular association with Rsu-1, and thus represent a regulatory mechanism for isoform and species-specific interactions of PINCH with Rsu-1.

Our structural and functional studies provide a basis for understanding why Rsu-1 is preferentially involved in the interaction with PINCH-1 at the downstream cluster of integrin-mediated adhesion complex. We showed here that the LRR domain of Rsu-1 is essential for the binding to PINCH-1 LIM5 domain and crucial for its localization of focal adhesion. More importantly, we found that Rsu-1 forms a robust tight association with the heterotrimer IPP complex that subsequently interacts with Kindlin-2, resulting in a tight heteropentamer complex (KIPPR complex). Our biophysical experiments suggest that the interaction between Rsu-1 and PINCH-1 substantially contributes to the stabilization the IPP-mediated focal adhesion assembly that is in agreement with previous genetic results (8,16). In support of this notion, we found that siRNA-mediated knockdown of Rsu-1 coincided with the reduction of PINCH-1 protein level, and the disruption of the Rsu-1-PINCH-1 interaction resulted in the reduction of focal adhesion formation and cell spreading. This notion of inter-molecular dependency is reminiscent to those in the formation of the IPP complex that each of the three proteins (ILK, PINCH, and Parvin) is crucial for the stability of each component and the localization to the sites of integrin adhesion complex (14). Notably, the IPP complex, along with Kindlin-2 has been implicated in the regulation of fundamental cellular processes (2,15). Hence, Rsu-1 significantly contributes to the stabilization of the KIPP-mediated focal adhesion assembly, thereby supporting the regulation of downstream pathways of integrin-associated signaling.

Previous studies demonstrated that the connection of Rsu-1 to PINCH-1 affected several downstream signaling pathways of MAPK (8,19,37). It remains to be determined whether the Rsu-1-PINCH-1 axis is involved in the regulation of oncogenic transformation (21). A previous study of affinity purification-mass spectrometry has identified that Raf-1 is involved in the IPP interaction network, suggesting an important link between the Rsu-1/IPP complex and the Ras/MAPK-mediated pathway (45). The LRR-containing proteins exhibit versatile structure frameworks that may implicate in various cellular functions (10). Previous genome-wide study of human LRR proteins has identified that Rsu-1 was an upregulated LRR protein with elevated expression in immune tissues (22). Thus, it remains to be determined whether Rsu-1 in concert with the IPP complex and other as-yet-uncharacterized molecules may play a role in the regulation of MAPK pathway. Deciphering molecular mechanisms responsible for the regulation of MAPK pathway by the Rsu-1 associated complex may cultivate further our understanding of how Rsu-1 is involved in regulating tumor development.

### Experimental Procedures

#### Antibodies and reagents

Rabbit polyclonal anti-Ras suppressor protein-1 (Rsu-1) primary antibody was purchased (Proteintech Group, Inc. and Thermo Fisher Scientific). Rabbit anti-HA and mouse anti-Myc epitope monoclonal antibodies were obtained from Cell Signaling Technology (Danvers, MA). Mouse monoclonal anti-pentahistidine antibody, mouse monoclonal anti-GST antibody, and anti-mouse and anti-rabbit secondary antibodies conjugated to horseradish peroxidase were from EMD Chemicals, Inc. (San Diego, CA). All chemicals and reagents were of analytical grade and purchased from Sigma-Aldrich unless otherwise specified.

#### Plasmids

The cDNA encoding human Rsu-1 (residues 1-277; Open Biosystems) was PCR-amplified and subcloned into pFastBac Dual (Life Technologies) with an engineered N-terminal hexahistidine-tag sequence followed by a thrombin cleavage site (designed as pFBDual-HT). For mammalian expression, Rsu-1 was subcloned into pCMV-HA (Clontech) to yield HA-tagged Rsu-1, into pCMV-Myc (Clontech) to yield Myc-tagged Rsu-1, and into pEGFP-c2 (Clontech) to yield GFP-fused Rsu-1. The bacterial expression plasmids for GST-fused various recombinant proteins were generated as previously described (38). Briefly, each gene encoding human PINCH-1 LIM4-5 (residues 189-325), LIM5 (residues 249-325), PINCH-2 LIM5L (residues 254-341; GenScript), PINCH-2 LIM5S (residues 254-330), mouse Raf-1 CR1 comprising Ras-binding domain and cysteine-rich domain (residues 54-188, DNASU) was PCR-amplified and subcloned into pGEX4T1 (GE Healthcare). The gene encoding the human HRAS (residues 1-166, DNASU) was PCR-amplified and subcloned into pET15b (Novagen). The amino acid substitutions or truncation in those expression plasmids were generated by site-directed mutagenesis with QuikChange Site-Directed Mutagenesis kit (Agilent Technologies) with appropriate primer sets. The tricistronic coexpression plasmid for the IPP complex was created as previously demonstrated (38). In brief, the cDNA of each human ILK (residues 1-452), α-parvin (residues 1-372), and PINCH-1 (residues 1-325) was PCR-amplified and subcloned into pET3aTr followed by co-expression vector of pST39 according to the inventor’s protocol (47). The bacterial expression plasmid for the Ulp1 cleavable hexahistidine and SUMO-tagged human kindlin-2 (residues 1-680) was created as previously demonstrated (40). All the DNA constructs were verified by sequencing analysis at Eurofins (Louisville, KY). Bacterial and mammalian DNA constructs were purified by QIAprep spin Miniprep kit (QIAGEN) and PureYield Plasmid Midiprep System (Promega), respectively.

### Generation of recombinant baculovirus

The pFBDual-HT plasmid containing Rsu-1 was transformed into MAX Efficiency DH10Bac *Escherichia coli* cells to generate recombinant bacmid DNA according to the manufacturer’s protocol (Bac-to-Bac Baculovirus Expression System, Thermo Fisher Scientific). The recombinant bacmid DNA was transfected into *Spodoptera frugiperda*-9 (Sf-9) insect cells with Cellfectin II reagent (Thermo Fisher Scientific) and the cells were incubated at 27°C in Sf-900 II SFM with penicillin and streptomycin in 6-well plate to generate the P1 baculoviral stock. The high-titer recombinant baculoviral stock was generated by amplification with infected Sf-9 cells in Sf-900 II SFM for three rounds and was used for large-scale protein expression. For producing recombinant Rsu-1 mutant proteins, each mutant plasmid was transposed into bacmid DNA in *E. coli* DH10Bac cells, and each recombinant bacmid DNA was purified and transfected into *Sf*-9 cells as for the wild type. The high-titer recombinant baculovirus of each mutant was amplified as for the wild-type.

### Expression and purification

The recombinant hexahistidine-tagged Rsu-1 was expressed in the baculovirus-infected *Sf*-9 insect cells grown at 27°C in ESF 921 serum-free culture medium with penicillin and streptomycin (Expression Systems). The cells were harvested at 72 hours post infection and lysed through freezing and thawing followed by a sonication in the buffer of 20 mM Tris, pH 7.5, 300 mM NaCl, 5% glycerol, 0.1% (v/v) NP-40, and an EDTA-free protease inhibitor cocktail (cOmplete) (Millipore Sigma). The lysates were clarified by centrifugation at 40,000 x g for 1 hour at 4°C. The hexahistidine-tagged Rsu-1 was purified from the lysate supernatant by Ni-affinity chromatography column equipped on an ÄKTA Purifier protein purification system (GE Healthcare). The fractions containing Rsu-1 were subjected to a thrombin cleavage to remove the hexahistidine-tag followed by an adjustment to a low salt buffer (20 mM Tris, pH 7, 30 mM NaCl). The thrombin cleavage was terminated by adding the above inhibitor cocktail. The Rsu-1 protein was further purified by cation exchange chromatography column with HiTrap SP HP followed by size exclusion chromatography column with Superdex 200 10/300 GL (all from GE Healthcare). Each Rsu-1 mutant protein (FRAA or R2W) was expressed in baculovirus-infected Sf-9 cells and purified as for the wild type.

The selenomethionine (SeMet)-substituted Rsu-1 protein was produced according to a protocol similar to previous study (48). In brief, *Sf*-9 cells at densities of 2 × 10^6^ cells/mL were grown in ESF 921 serum-free medium and infected with the recombinant baculovirus. After an optimized period of infection (between 16 to 24 hours), the cells were harvested by centrifugation at 500 x g for 10 min and resuspended in ESF 921 Delta Series Methionine Deficient medium (Expression Systems). The cells were incubated in the methionine-free medium for 4 hours to deplete the intracellular pool of methionine (48) and exchanged by centrifugation as the above for the labeling medium of ESF 921 Delta Series Methionine Deficient medium supplemented with 50 μg/mL L-SeMet (Millipore Sigma). The cells were grown in the labeling medium for 72 hours and harvested by the centrifugation as the above. The SeMet-substituted Rsu-1 protein was purified from the lysate of the infected *Sf*-9 cells as for the native protein described in the above.

The recombinant proteins of various GST-fused PINCH proteins were expressed and purified as described below. In brief, *E. coli* strain Rosetta 2 (DE3) cells harboring each bacterial expression plasmid were grown at 20°C in LB medium for 16-20 hours after induction with 0.2 mM IPTG. Cells were harvested by centrifugation at 5,000 x g for 20 min and lysed by sonication in 20 mM Tris, pH 7.5, 150 mM NaCl, 5% glycerol, 50 μM zinc acetate, and EDTA-free protease inhibitor cocktail (cOmplete). Each lysate was cleared by centrifugation at 40,000 x g for 1 hour and loaded onto GSTrap affinity chromatography column (GE Healthcare) equilibrated in the same buffer. Each bound protein was eluted in 25 mM reduced glutathione in the buffer. Cleavage from GST upon necessary was carried out with α-thrombin, and the reaction mixtures were subjected to buffer exchange to a low salt buffer consisting 20 mM Tris, pH 7, 30 mM NaCl. Each cleaved PINCH protein was loaded onto HiTrap SP cation exchange chromatography and eluted using a linear NaCl gradient. Each PINCH protein was further purified by size exclusion chromatography on a HiLoad 16/60 Superdex 75 column (GE Healthcare) equilibrated in 20 mM Tris, pH 7.5, 150 mM NaCl, and 0.02% NaN_3_.

The hexahistidine-tagged IPP heterotrimer complex was expressed in *E. coli* and purified from the bacterial cell lysate by Ni-affinity chromatography column followed by HiLoad 16/60 Superdex 200 size exclusion and HiTrap SP chromatography columns, as previously demonstrated (38). The hexahistidine and SUMO-tagged kindlin-2 was expressed in *E. coli* and purified from the bacterial cell lysate by Ni-affinity chromatography column followed by HiTrap Q and HiLoad 16/60 Superdex 200 chromatography columns. The SUMO-tag was cleaved by Ulp1 digestion, as previously demonstrated (40). The GST-fused Raf-1 CR1 domain (RBD and CRD) was expressed in *E. coli* as the above and purified from the bacterial cell lysate by GSTrap affinity chromatography followed by HiLoad 16/60 Superdex 200 size exclusion chromatography. The hexahistidine-tagged HRAS (G12V) was expressed in *E. coli* and purified from the bacterial cell lysate by Ni-affinity chromatography column followed by HiLoad 16/60 Superdex 200 size exclusion chromatography column. The N-terminal hexahistidine-tag was removed by a thrombin cleavage.

### Complex formation

The protein complexes between Rsu-1 and the PINCH-1 LIM4-5 domains for crystallization experiments were prepared by incubating each purified protein in a 1:1 molar ratio by a rotor at 4°C for at least 2 hours. The protein complex mixtures were loaded onto a size exclusion chromatography column of either Superdex 200 10/300 GL (GE Healthcare) or Superdex 75 Increase 10/300 (GE Healthcare) pre-equilibrated in a buffer consisting of 20 mM Tris, pH 7.5, and 150 mM NaCl. The major peak fractions containing the target protein complex were pooled and concentrated with Vivaspin 20 (MWCO 10K) centrifugal concentrator for crystallization experiments. The protein concentration was quantitated by measuring absorbance at 280 nm with NanoDrop 2000c Spectrophotometer from Thermo Fisher Scientific (Waltham, MA).

### Crystallization and data collection

Initial crystallization screens were carried out by sitting-drop vapor diffusion method with Gryphon (Art Robbins Instruments). All the crystallization screening plates were incubated at 23°C. The best native crystals of Rsu-1 protein were obtained by mixing 1 μL of Rsu-1 (30 mg/mL) with the equal volume of the reservoir solution containing 0.1 M Tris, pH 8.5, 0.8 M LiCl, and 30% PEG4000. The SeMet-substituted Rsu-1 protein was crystallized under conditions nearly identical to the native Rsu-1. The crystals of the complex (19.3 mg/mL) between Rsu-1 and a tandem repeat of LIM4-5 domains of PINCH-1 were grown in the reservoir solution consisting 0.1 M Hepes, pH 7.5 and 11% PEG8000. Those crystals were soaked in cryopreservation in 25% glycerol in the reservoir solutions and stored in a liquid nitrogen tank until data collection. Crystallographic data collection experiments were carried out at 100K at the Advanced Photon Source Structural Biology Center 19-BM beamline using a wavelength (λ=0.97919 Å) for the native crystals of Rsu-1. For the SeMet-substituted crystal, an X-ray fluorescence scan was carried out and the resultant data were analyzed to determine the presence of selenium atoms and location of the absorption edge for the crystal. A single-wavelength anomalous diffraction data set for a single SeMet-substituted crystal was collected at the selenium peak wavelength (λ=0.97941 Å). Those crystallographic data were integrated and scaled with the program HKL3000 (49).

### Structure determination and refinement

The structure of Rsu-1 was determined by a single-wavelength anomalous dispersion (SAD) phasing method with data collected from a single crystal of selenomethionine-substituted protein. Five selenium atoms were located with the program SHELXD (50) in the HKL2MAP interface (51) and utilized for phasing and density modification with the program SHELXE (52) that resulted in an estimated mean figure-of-merit of 0.715 and pseudo-free correlation coefficient of 74.93%. The handedness of the solution was determined by inspecting and analyzing the resultant experimental maps and their connectivity. The experimental SAD map with correct hand showed clear and interpretable continuous density, revealing secondary structures corresponding to the most part of the LRR domain of Rsu-1. The initial model (polyalanine) was generated with SHELXE (52) and further constructed with an aid of automated model building using the program ARP/wARP (53). Iterative rounds of manual model building and refinement were carried out with the programs COOT (54) and REFMAC5 (55). The resultant model was then utilized to solve the structure of Rsu-1 in native methionine (unlabeled) form using the program MOLREP (56) in the CCP4 package (57). Manual rebuilding was carried out with COOT (54), and the structure of Rsu-1 in the native form was refined by the programs REFMAC5 (55) and PHENIX (58). Addition of water molecules was carried out with COOT (54).

The crystal structure of the complex between Rsu-1 and a tandem repeat of LIM4-5 domains of PINCH-1 was determined by molecular replacement with the program PHASER (59) using the atomic coordinates of Rsu-1 as a search model. Initial molecular replacement solution was obtained from data of crystals that were grown in initial crystallization conditions (0.1 M sodium phosphate, pH 6.5, 12 % PEG 8,000, 0.2 M sodium chloride). The initial crystals of the complex diffracted to a resolution of 3.05-Å. The structure solution resulted in interpretable density maps for Rsu-1, the LIM5 domain of PINCH-1, and their binding interface but the quality of the map for the LIM4 domain was not sufficient probably owing to a disordered LIM4. The LIM5 domain was manually built on the basis of interpretation of the electron density map (2*F*_o_-*F*_c_); however, the map quality for the LIM4 domain was not improved during model building and refinement using this dataset. Meantime, a new dataset of the crystals of the complex that were grown in a distinct crystallization condition was available. Despite a moderate diffraction to a resolution at 3.35-Å from the new crystals, the molecular replacement using the atomic coordinates of the partially refined complex between Rsu-1 and LIM5 as templates resulted in an improved interpretable electron density map for the LIM4 domain. Iterative model building for LIM4 domain was carried out based on the new density map from the new dataset. The structure of the complex between Rsu-1 and a tandem repeat of the LIM4-5 domains of PINCH-1 was further refined with REFMAC5 (55) and PHENIX (58). The stereochemistry of the final coordinates was assessed by the programs PROCHECK (60) and MOLPROBILY (61). Crystallographic data collection, phasing, and refinement statistics are summarized in Table 2.

### Structural, modeling, and sequence analysis

The structural superpositions were carried out with the secondary structure matching (SSM) algorithm supplemented in COOT. The intermolecular residue contacts were analyzed with PISA (33) and CONTACT from the CCP4 suite (57). A homology model of the PINCH-2 LIM5 domain was constructed with the SWISS-MODEL (62) using the atomic coordinates of the PINCH-1 LIM5 domain taken from the bound form with Rsu-1 (this study) as a template. The resultant homology model that lacks the C-terminal extended 11-residue was superposed onto the PINCH-1 LIM5 domain bound to Rsu-1 to generate a comparative structural model of the complex between Rsu-1 and PINCH-2 LIM5S. The secondary structure assignments were carried out with the program STRIDE (63). Electrostatic surface potential maps were calculated with the program APBS (64). Buried surface areas in the interface structures were calculated with the program CNS (65) using a probe radius of 1.4 Å. Shape complementatiry at the protein-protein interfaces were analyzed by measuring the shape correlation statistics (Sc) (31). The sequence conservation and mapping the scores on the structure were analyzed with the program ConSurf (66) using default parameters and the Bayesian method. Multiple sequence alignments were carried out by CLUSTAL Omega (67) and rendered with the program ESPript (68). Structural figures were generated with the program PYMOL (http://pymol.org).

### Interaction analysis by size exclusion chromatography

The protein interaction experiments were carried out on either a Superdex 75 Increase 10/300 GL, a Superdex 200 10/300 GL, or a Superose 6 10/300 GL column (all from GE Healthcare). Those columns were pre-equilibrated in a buffer A consisting of 20 mM Tris, pH 7.5, 150 mM NaCl or a buffer B consisting of 20 mM Hepes, pH 7.5, 150 mM NaCl, and 0.2 mM TCEP, and calibrated with gel filtration standard proteins comprising thyroglobulin (670 K), γ-globulin (158K), ovalbumin (44K), myoglobin (17K), and vitamin B12 (1.35K) (BIO-RAD). For the binary complex formation between Rsu-1 and each PINCH-1/-2 LIM5 protein, approximately 200 μg of each purified LIM5 (PINCH-1 LIM5 wild type or PINCH-2 LIM5S) was mixed with equimolar amount of purified Rsu-1 (wild type or loss-of-binding mutants such as FRAA or R2W), incubated at 4°C by a rotor for at least two hours, and run onto the column. Control experiment for each unbound protein was analyzed in the same method. For the complex formation of the heteropentamer complex (KIPPR), 1.35 mg of purified IPP heterotrimer protein was mixed with approximately 400 μg of purified Kindlin-2 and 325 μg of purified Rsu-1 in a buffer C consisting of 20 mM Tris, pH 7.5, 150 mM NaCl supplemented with 5% glycerol and 0.2 mM TCEP. The mixture of those proteins was incubated at 4°C by a rotor for at least 2 hours and run onto a Superose 6 column, as described the above. The major peak fractions containing the KIPPR heteropentamer complex that were eluted from 12.89 ml to 14.89 ml were collected and concentrated to a minimal volume with a Vivaspin 20 (MWCO 10 K) centrifugal concentrator. The concentrated sample was then filtered and re-loaded onto the same column, and the stable heteropentamer complex was eluted at the same fractionated position as a single peak on the column. Control experiment for each unbound protein was carried out on the column in the same way.

### Biolayer interferometry

Real-time binding interactions between Rsu-1 and PINCH-1/-2 proteins were measured by biolayer interferometry with a single-channel BLItz instrument (Pall FortéBio). Prior to the measurements, the anti-GST biosensors were hydrated for at least 10 minutes in a BLItz buffer consisting of 20 mM Tris, pH 7.5, 150 mM NaCl, 1 mg/mL BSA, and 0.05% (v/v) Tween-20 in a 96-well black flat-bottom microplate (Greiner Bio-One) in an adaptor tray (Forté Bio). Each experiment comprised five steps: initial baseline (30 seconds), loading (120 seconds), baseline (30 seconds), association (120 seconds), and dissociation (120 seconds). Loading was performed using each GST-fused protein or GST at 0.05 mg/ml diluted in the BLItz buffer, and binding interactions were measured during association with Rsu-1 WT or mutant proteins at various concentrations diluted in the same buffer. Each buffer (250 μL) was maintained in a 0.5 mL black microcentrifuge tube (Cole-Parmer) and exchanged immediately before each dissociation phase. Binding signals were measured in real-time in nanometers (nm) as a function of time (seconds), and the binding sensorgrams were normalized by subtracting a reference run with the BLItz buffer. Data acquisition and analysis for kinetic measurements were carried out using the program BLItz Pro (Forté Bio). The binding affinity (KD) was obtained by 1:1 global fitting model in the BLItz Pro software. The binding experiments were repeated for at least three-independent measurements.

### Protein thermal shift assay

Experiments were carried out using a QuantStudio 5 Real-Time PCR System from Thermo Fisher Scientific. Proteins were buffer-exchanged in 20 mM Tris, pH 7.5, 150 mM NaCl and prepared in a MicroAmp Optical 96-well plate from Thermo Fisher Scientific at a final concentration of 2 μM in 20 μL reaction volume that contained a Protein Thermal Shift Dye (Thermo Fisher Scientific) as a fluorescence probe at a 1X concentration. The temperature was raised in steps at a ramp rate of 0.05°C per second from 25°C to 99°C. The excitation and emission filters were set to x4 (580±10 nm) and m4 (623±14 nm), respectively. The thermal melting measurements were performed for three independent experiments in four replicates of each reaction. For the reference wells, no protein control (NPC) was prepared in the reaction mixture consisting buffer and dye without protein. The melting temperature (T_m_) was determined by fitting the melt curve to the Boltzmann equation using the Protein Thermal Shift Software version 1.0 from Thermo Fisher Scientific.

### Cell culture and transfection

HeLa-2F5 cells (CRISPR generated PINCH-1 deficient cell line) were maintained in Dulbecco’s Modified Eagle’s Medium supplemented with 10% fetal bovine serum, as previously demonstrated (38). MCF10A cells (ATCC CRL-10317) were maintained in Mammary Epithelial Cell Growth Medium (Lonza#CC-3150) supplied with 100 ng/ml cholera toxin. All cells were kept at 37°C in incubator with 5% CO_2_. Endogenous Rsu-1 proteins were transiently knockdown by siRNA (RSU1_s02) using Lipofectamine RNAiMAX Transfection Reagent (Fisher) for 48 hours. Rsu-1 knockdown was followed by transient expression of HA-Rsu-1 constructs in MCF10A cells, using jetOPTIMUS (Polyplus). Co-expression of EGFP-PINCH-1 (38) and Rsu-1 constructs in HeLa-2F5 were performed using PEI reagent.

### Western blot

Cells were lysed in lysis buffer (50mM Tris-HCl, pH 6.8, 1% SDS) and the protein lysate was quantitated with Pierce BCA protein assay kit (Fisher). Quantitated lysates were diluted in Laemmli buffer containing 62.5 mM Tris-HCl, pH 7.4, 2% SDS, 5% 2-mercaptoethanol, and 10% glycerol and subjected to SDS-PAGE. Proteins were then transferred to PVDF and probed with primary antibodies (anti-GFP XP, Cell Signaling; anti-HA, Cell Signaling; anti-PINCH, BD Bioscience; anti-RSU1, Invitrogen; anti-GAPDH, Cell Signaling) which was followed by HRP conjugated secondary antibody (anti-mouse HRP, Cell Signaling; anti-rabbit HRP, Cell Signaling).

### Cell spreading assays

After DNA transfection, MCF10A cells or HeLa cells were seeded to fibronectin coated coverslips (10 µg/cm_2_) for 2 hours spreading at 37°C. Adherent cells were fixed with 4% formaldehyde and stained with anti-vinculin antibody (Sigma) or anti-GFP antibody (abcam) and anti-HA antibody (Cell Signaling) followed by goat anti-mouse antibody Alexa 488-conjugated (abcam) or goat anti-chicken antibody Alexa 488-conjugated (abcam) and goat anti-rabbit Alexa 568-conjugated (abcam). Coverslips were then mounted with Prolong Diamond Antifade Reagent with DAPI (Fisher) overnight and visualized with Leica TCS-SP5 II upright confocal microscope (Leica Microsystems, GmbH, Wetzlar, Germany). Images were followed with ImagePro 10 software processing for spreading area and FA size/number quantification.

### Statistical Analysis

All data were compared using SigmaPlot 10.0 software by Mann-Whitney Rank Sum Test due to failed normality tests and equal variance tests. Differences were considered to be significant when P< 0.05.

## Data Availability

The atomic coordinates of Rsu-1 (free form) and its complex with the PINCH-1 LIM4-5 domains have been deposited in the Protein Data Bank (accession codes: XXXX and YYYY, respectively).

## Supporting Information

This paper is associated with 4 supporting figures and 1 supporting table.

## Acknowledgments

The crystal structures in this report are derived from works performed at Structural Biology Center (SBC) beamline 19-BM at the Advanced Photon Source in the Argonne National Laboratory. The Argonne National Laboratory is operated by University of Chicago Argonne, LLC, for the US Department of Energy, Office of Biological and Environmental Research under contract DE-AC02-06CH11357. We thank to Song Tan for the use of polycistronic coexpression system.

## Funding and Additional Information

This work was supported by the National Institutes of Health (5R01HL058758: J.Q.). The content is solely the responsibility of the authors and does not necessarily represent the official views of the National Institutes of Health.

